# emb2dis: a novel protein disorder prediction tool based on ResNets, dilated convolutions & protein language models

**DOI:** 10.64898/2026.03.30.715414

**Authors:** S.A. Duarte, M. Mehdiabadi, L.A. Bugnon, M.C. Aspromonte, D. Piovesan, D.H. Milone, S.C.E. Tosatto, G. Stegmayer

## Abstract

Intrinsically disordered proteins (IDPs) play an important role in a wide range of biological functions and are linked to several diseases. Due to technical difficulties and the high cost of experimental determination of disorder in proteins, combined with the exponential increase of unannotated protein sequences, the development of computational methods for disorder prediction became an active area of research in the last few decades. In this work, we present emb2dis, a deep learning model that uses protein language models (pLMs) to predict disorder from sequence. The emb2dis tool is a pre-trained model that receives as input a protein sequence, calculates its pLM embedding and passes it to a deep learning model. In contrast to existing approaches, emb2dis integrates informative sequence representations with a novel architecture that combines residual networks (ResNets) and dilated convolutions. This design effectively enlarges the receptive field of the convolution operation, enabling the model to better capture an extended context of each amino acid. At the output, emb2dis assigns a disorder propensity score to each residue in the sequence. The model was evaluated on datasets from the latest CAID3 blind benchmark for disorder prediction, where it achieved first place in the Disorder-PDB category, exhibiting strong performance with high AUC and *F*_*max*_ scores. Additionally, it ranked among the top ten methods on the Disorder-NOX dataset. We provide a freely available web-demo for emb2dis and a source code repository for local installation.

**Weblink for the tool:** https://sinc.unl.edu.ar/web-demo/emb2dis/

The importance of the emb2dis tool is that it provides a new deep learning approach and significant improvements in the prediction of protein disorder, with a simple web interface and graphical output detailing per-residue disorder.

## Introduction

Intrinsically disordered proteins (IDPs) and intrinsically disordered regions (IDRs) lack a well-defined three-dimensional structure, resulting in high conformational flexibility under physiological conditions (Dunker et al., 2001; van der Lee et al., 2014). At the sequence level, IDRs are characterized by amino acid compositions that favor dynamic polypeptide chains unable to adopt a stable tertiary fold, often due to a low proportion of hydrophobic residues required to form a compact hydrophobic core. Despite the absence of a stable 3D structure, IDPs and IDRs are essential for cellular function. Their structural plasticity enables them to participate in a wide range of biological processes, such as transcription, translation, signaling, cell division, and cellular homeostasis (Holehouse & Kragelund, 2023). IDRs can vary in length, ranging from short segments of a few residues to extended regions spanning hundreds or even thousands of amino acids.

Although IDPs and IDRs can be investigated and characterized experimentally, their intrinsic flexibility poses significant challenges for both experimental characterization and computational predictions. In the last few years, significant progress has been made in the development of computational tools for predicting disorder directly from protein sequences (Erdős & Dosztányi, 2024; Uversky & Kurgan, 2023, Han et al., 2023). Benchmarking initiatives such as the Critical Assessment of Intrinsic Disorder (CAID) challenge have provided a rigorous framework for comparing predictors and highlighting remaining gaps in performance, particularly in low-confidence or ambiguous regions (Mehdiabadi et al., 2026; Del Conte et al., 2023).

In recent years, one promising direction for improving current disorder prediction models has emerged from the field of deep learning (DL): the use of protein language models (pLMs), which are deep neural networks pre-trained on millions of sequences using self-supervised learning. These models generate high-dimensional representations that encode sequence-level information relevant to structure and function (Rao et al., 2020; Vitale et al., 2024). A pLM can capture some aspects of the language used to write protein primary sequences (Weissenow & Rost, 2025). Early large language models in natural language processing were trained by masking out a few words in a sentence and learning to predict them from the context provided by the other words, a process known as pre-training. In a pLM, words are replaced by residues, and the model is trained to predict masked segments of each input sequence. A pLM generally adopts the Transformer architecture, which forms an internal representation of the residues and their contexts (Vaswani et al., 2017). After pre-training, the output of the last encoder layer can be used to generate a compact numerical vector for each protein residue, known as embedding, which can be used for several downstream tasks such as the prediction of protein structure and function (Wang et al., 2025; Fan et al., 2025; Littmann et al., 2021).

Several pre-trained pLMs have emerged over the last few years (Yang et al., 2018; Detlefsen et al., 2022; Tran et al., 2023) that leverage the vast amount of unannotated protein sequence data available. Recent reviews (Fenoy et al., 2022; Vitale et al., 2024) experimentally benchmarked several protein sequence representation learning methods, indicating Evolutionary Scale Modeling (ESM) (Rives et al., 2021) and ProtT5 (Elnaggar et al., 2022) as the best methods for the downstream tasks evaluated. Other studies have shown that pLM embeddings can enhance performance in a wide range of predictive tasks, including disorder prediction (Heinzinger et al., 2024). Building on these advances, this work focuses on developing a DL model for disorder prediction based on pLM embeddings, named emb2dis. The model proposes a novel DL architecture based on ResNets and dilated convolutions. This approach was trained and evaluated across different datasets to assess its ability to identify disordered regions in protein sequences. A web-demo tool is provided at https://sinc.unl.edu.ar/web-demo/emb2dis/ and the repository including source code can be found in https://github.com/sinc-lab/emb2dis.git.

### The emb2dis disorder prediction model

The model proposed here is a convolutional neural network designed for sequence-level classification, as shown in Figure 1. For each protein, the corresponding full sequence of embeddings is obtained using a protein language model (pLM). The input to the model consists of windows of fixed length *W*, extracted from the full-protein sequence embeddings of length *L*. At test time, the window *W* is slid along the full sequence with a step size of 1 and an overlap of *W*-1, to obtain a per-aminoacid prediction. The emb2dis architecture includes an initial convolutional layer, followed by a stack of residual networks (ResNet) (He et al., 2016) with dilated convolutions (Yu et al., 2017) and bottleneck layers (Koh et al., 2020). The dilated convolutions are a variant of traditional convolutions that introduce gaps (dilations) between kernel elements, increasing the receptive field of the convolution operation by looking at larger areas than a normal convolution. This helps capture more global context from the input amino acids without increasing the number of parameters or the filter size. Additionally, dilated convolutions are more computationally efficient. The extracted features are then fed into an adaptive max pooling layer, which integrates information from the *F* features across the *W* residues of the window. Then, a dropout layer (Srivastava et al., 2014) is included for regularization in training, and a fully connected layer provides the model output. At this output, for each residue, there are two possible classes: structured or disordered.

**Figure 1.**
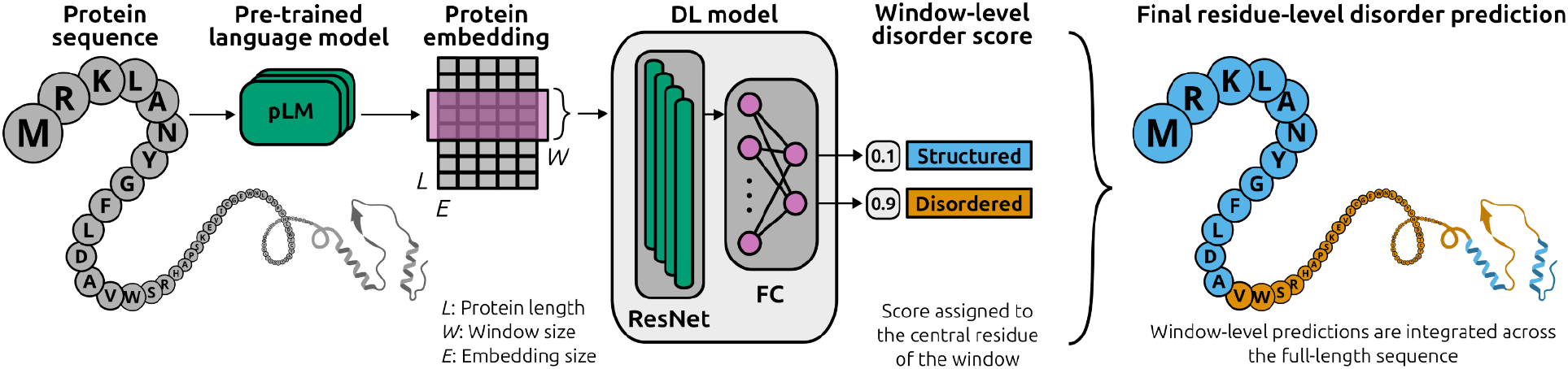
The emb2dis model architecture. For each input protein, a sequence of pLM embeddings is obtained. The model has a convolutional layer, a stack of residual networks (ResNet) with dilated convolutions and bottleneck layers. A fully connected layer provides the model output for each residue: structured or disordered. In test time, a window is slid along the input sequence to obtain a per-aminoacid prediction.

The model was optimized using the Adam algorithm (Kingma & Ba, 2017) with weight decay, and the loss function used was the standard cross-entropy. Two pLMs were used for embedding extraction: ESM (Rives et al., 2021), including ESM2 and ESMC; and ProtT5 (Elnaggar et al., 2022). ESM2 is an encoder-only transformer language model for proteins with up to 15 billion parameters, making it the largest pLM to date. ESMC for Cambrian is a replacement of the previous ESM models that provides major improvements in capability and efficiency with a much lower number of parameters. ProtTrans ProtT5 has an encoder-decoder transformer architecture. All of these pLMs have demonstrated remarkable potential in capturing structural and functional properties of proteins through pretraining on large-scale protein sequence data (Saadat & Fellay, 2025). In the case of ESM2, the embedding dimension is *E*=1,280 and for ESMC 600m is *E*=1,152, while for ProtT5, the embedding dimension is *E*=1,024. The embeddings are obtained per-residue. The main architectural and training hyperparameters were selected through optimization: learning rate (*lr*) (from 1e-2 to 1e-6), *W* length (from 10 to 100, with step size 5), number of 1D filters (from 50 to 400, with step size 50), kernel size (from 3 to 11), number of ResNet blocks (from 1 to 4) and dropout probability (in the 0.00 to 0.60 range, with step 0.10). The optimization process involved a total of 250 experiments with the training and validation sets (strictly excluding proteins in the test set). The first 50 trials used random sampling to ensure broad coverage of the search space. The remaining trials were guided by the tree-structured parzen estimator algorithm (Bergstra et al., 2011), which aimed to maximize the area under the ROC curve (AUC) on the validation set. After completing the hyperparameter optimization, the best-performing configuration was selected based on the highest AUC on the validation set. For emb2dis-ESM2, the final parameters are *lr*=1.2e-04, *W*=30, 400 filters, 2 ResNet blocks, kernel size of 11, and 0.20 for dropout rate. For emb2dis-ProtT5 instead, the *lr* was 2e-05, *W*=35, 200 filters, and 1 ResNet block, without dropout. For emb2dis-ESMC 600m the *lr* was 2e-06, 400 filters, kernel size of 9, and 0.30 for dropout rate.

### Datasets and annotations

IDR annotations for developing our model were extracted from the DisProt database (v9.5 release 2023_12), a resource that collects high-quality IDP/IDR annotations based on literature evidence (Nugnes et al., 2026). The training data were generated following the reference definition of Disorder-PDB in the CAID challenge, where observed residues in experimentally determined PDB structures are used as negative labels. Residues annotated as disordered in DisProt but structured in the PDB were considered as disordered. Residues without any structural or disorder annotation were excluded from the analysis and were not used for training. Only proteins with less than 2,000 amino acids were used to create the training dataset, due to computational resources and pLMs limitations. Redundancy in the dataset was eliminated using CD-HIT (Fu et al., 2012), with a maximum sequence identity threshold of 40%. After applying all filtering criteria, the final dataset included 2,246 unique protein sequences. We generated embeddings as in Vitale et al. (2024), obtaining protein representation using three pLMs: ESM2, ESMc 600m, and ProtT5.

This dataset was split into two partitions: a training set (90%) and an internal validation set (10%). The generalization capability of the model was monitored during training using the validation set. At validation, the AUC and *F*_*max*_ were very close to those obtained in training, indicating a robust performance in identifying disordered regions without overfitting. For example, for emb2dis-ESM2, the AUC and *F*_*max*_in the train set were 0.941 and 0.880, respectively; while in validation, these metrics were 0.921 and 0.857. In the case of emb2dis-ProtT5, the AUC and *F*_*max*_ in training were 0.976 and 0.858, respectively, while in validation, they were 0.921 and 0.923, respectively. The same trend was verified for emb2dis-ESMC.

For testing, this study used the benchmark datasets already provided by the CAID3 challenge (Mehdiabadi et al., 2026). Specifically, two datasets were evaluated: Disorder-PDB and Disorder-NOX. In the Disorder-PDB dataset, all disorder-annotated residues in DisProt were considered positive for the benchmark, and negative residues were constrained to observed regions in the PDB. The challenge authors state that this dataset is conservative and more reliable than the other datasets because it excludes uncertain residues without structural or disorder annotation. In the Disorder-NOX dataset, the challenge clarifies that all disorder-annotated residues in DisProt, excluding X-ray missing residues, were considered positive, and the rest were negative. The missing residues were excluded from the evaluation using PDB structural information (Mehdiabadi et al., 2026). Additionally, for performance comparison, we downloaded the predictions from state-of-the-art models submitted to CAID3 from the official website and used them for comparative analysis. The model was evaluated using the assessment scripts of the official CAID repository (https://github.com/BioComputingUP/CAID). The same classification metrics as the CAID3 challenge were calculated: AUC, Average Precision Score (APS) and *F*_*max*_.

## Results

### Performance comparison with state-of-the-art methods

Table 1 reports the results of the three variants of emb2dis on the Disorder-PDB benchmark dataset of the CAID3 challenge. The table shows the top-10 methods for this benchmark dataset. Evaluation showed that emb2dis-ESM2 achieved the highest AUC and *F*_*max*_ scores compared to the other state-of-the-art methods that participated in the challenge. Remarkably, all the other variants of emb2dis are also in the top-10 table, with emb2dis-ESMc in 3rd place and emb2di-ProtT5 in 8th place, according to AUC. In terms of APS, however, the best model is em2dis-ESMC.

**Table 1.** Performance of the top-10 predictors on the Disorder-PDB benchmark dataset. The best score for each metric is indicated in bold.

| Model | AUC | APS | $F_{max}$ |
| --- | --- | --- | --- |
| **emb2dis-ESM2** | **0.956** | 0.927 | **0.860** |
| PUNCH2 (Meng & Pollastri, 2025) | 0.955 | 0.929 | 0.859 |
| **emb2dis-ESMC** | 0.953 | **0.931** | 0.859 |
| PUNCH2-Light (Meng & Pollastri, 2025) | 0.953 | 0.925 | 0.855 |
| AlphaFold-rsa (Piovesan et al., 2022) | 0.950 | 0.921 | 0.858 |
| SPOT-Disorder2 (Hanson et al., 2020) | 0.949 | 0.920 | 0.840 |
| AlphaFold3-rsa | 0.948 | 0.913 | 0.856 |
| **emb2dis-ProtT5** | 0.947 | 0.922 | 0.850 |
| LMDisorder (Song et al., 2023) | 0.937 | 0.898 | 0.825 |
| ESMDisPred-2PDB (Kabir et al., 2026) | 0.937 | 0.893 | 0.837 |

Figure 2 presents the A) ROC curve and B) precision-recall curve for all evaluated predictors on the Disorder-PDB benchmark dataset. On the ROC curve, emb2dis-ESM2 is the best-performing model with an AUC of 0.956. It has to be mentioned that actually all methods are very close to each other. The precision-recall curve shows that em2dis-ESMC is the best model, with an APS of 0.931, followed by emb2dis-ESM2 in 3rd place and emb2dis-ProtT5 in 5th place. According to this curve and the APS score, again, all three variants of emb2dis are among the top-10 best-performing methods in this benchmark dataset.

**Figure 2.**
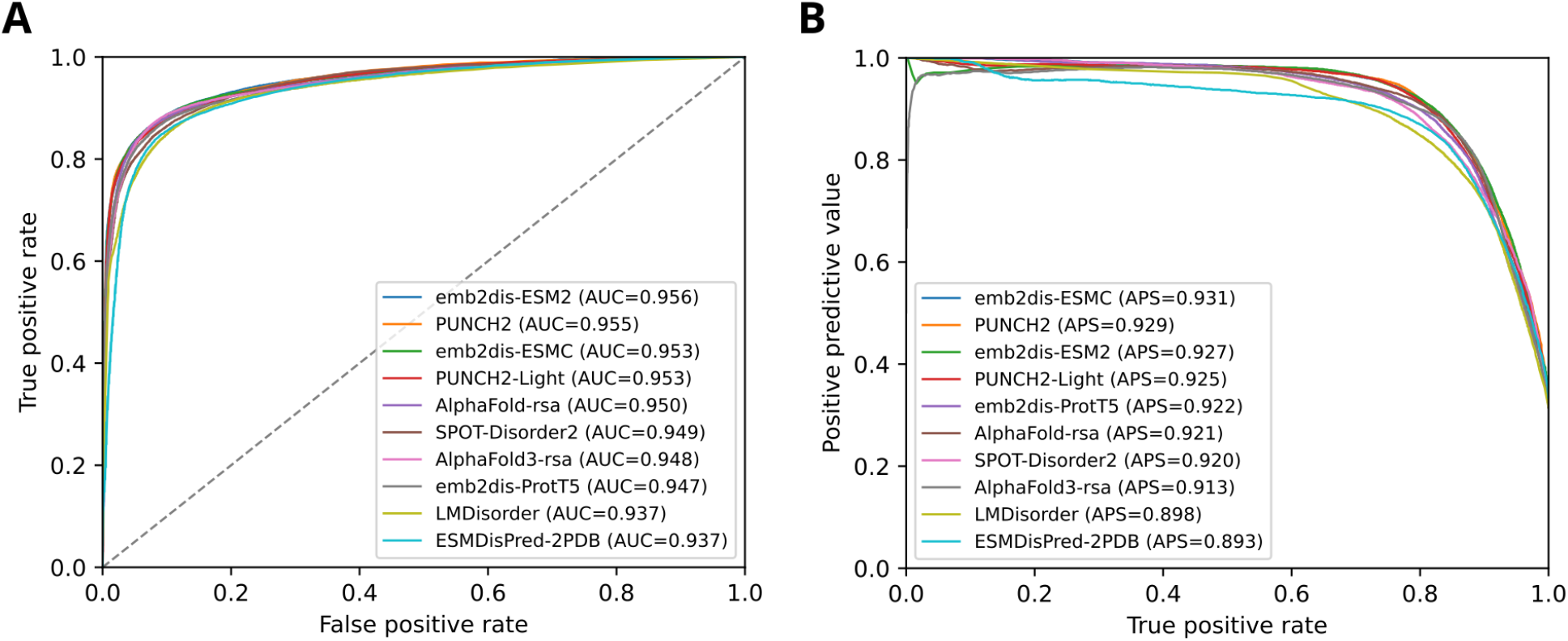
A) ROC curve with false positive rate (x-axis) and true positive rate (y-axis), for the Disorder-PDB benchmark dataset of CAID3. B) Precision-recall curves on the same benchmark dataset.

Table 2 shows the comparative results of the top-10 prediction methods for the Disorder-NOX dataset. Now, for the emb2dis models, some variability in the metrics is observed with respect to the previous dataset, which is expected due to differences in the annotation criteria and dataset composition. For example, the Disorder-PDB dataset includes only residues with known annotations: residues observed in PDB structures are labeled as structured (negatives), while those annotated as disordered in DisProt are labeled as positives. This is the same strategy adopted in our training dataset, which excludes ambiguous or unannotated residues. In contrast, the Disorder-NOX dataset excludes annotations of missing residues and labels any residue not annotated as disordered as structured. Here, the model emb2dis-ESM2 ranked 6th, achieving competitive performance within the top 10 methods, even under more challenging conditions. The model emb2dis-ESMC ranked 9th. It should be noted that emb2dis-ProtT5 ranked 13th. In spite of the fact that emb2dis was not first in this more challenging dataset, it is notably the only model that is part of the top-10 results in both benchmark datasets. None of the other competitors’ methods are among the best predictors in both datasets simultaneously.

**Table 2.** Performance of the top-10 predictors on the Disorder-NOX benchmark dataset. The best score for each metric is indicated in bold.

| Model | AUC | APS | $F_{max}$ |
| --- | --- | --- | --- |
| ESMDisPred-2PDB (Kabir et al., 2026) | **0.885** | **0.754** | **0.750** |
| ESMDisPred-1 (Kabir & Hoque, 2024) | 0.876 | 0.745 | 0.720 |
| ESMDisPred-2 (Kabir & Hoque, 2024) | 0.872 | 0.743 | 0.725 |
| DisoFLAG-IDR (Pang & Liu, 2024) | 0.868 | 0.714 | 0.671 |
| DisorderUnetLM (Kotowski et al., 2025) | 0.862 | 0.702 | 0.647 |
| **emb2dis-ESM2** | 0.861 | 0.630 | 0.667 |
| flDPnn3a (Wang et al., 2024) | 0.860 | 0.700 | 0.668 |
| UdonPred-combined (Schlensok et al., 2026) | 0.854 | 0.668 | 0.671 |
| **emb2dis-ESMC** | 0.851 | 0.649 | 0.660 |
| DisPredict3 (Kabir & Hoque, 2024) | 0.850 | 0.667 | 0.658 |

### Examples of protein disorder predictions with emb2dis

The analysis of model predictions along protein sequences offers a more detailed view of emb2dis performance. We will show examples that demonstrate the model’s ability to capture both the location and extent of disordered regions. Figure 3 shows the emb2dis-ESM2 prediction of the protein P10912, the human growth hormone receptor (DisProt ID: DP00033), which was part of the training set. The emb2dis disorder prediction curve is in orange, with the ground truth disorder indicated in an orange background and the ordered region in light blue. At the bottom, the pLDDT score for AlphaFold2 (AF2) of this protein is shown, along with the annotations it receives from different databases (DisProt, MobiDB, and UniProt). This protein is a member of the class I cytokine receptor family, which functions as a homodimer and plays a fundamental role in regulating postnatal growth and development. It consists of a conserved extracellular ligand-binding domain (residues 19-264), a single-pass transmembrane alpha-helix (265-288), and a cytoplasmic domain (289-638) that is largely intrinsically disordered and mediates downstream signaling (Haxholm et al., 2015). Most of these structural features are captured by our model, which correctly identifies ordered regions in the extracellular domain and the disorder across the intracellular region. Notably, a decrease in the emb2dis score is observed at the transmembrane segment, consistent with its alpha-helical secondary structure. It can be noticed that the predicted disorder by emb2dis-ESM2 is consistent with the curated annotations of DisProt. Additionally, emb2dis-ESM2 indicates a region at the beginning of the sequence that is not labeled in DisProt and might be disordered. Indeed, it is marked as missing residues in MobiDB and has a low pLDDT score, which is usually associated with disorder. Overall, even though our model does not require any structural information as input, the emb2dis score shows a strong correlation with the AF2 pLDDT confidence score.

**Figure 3.**
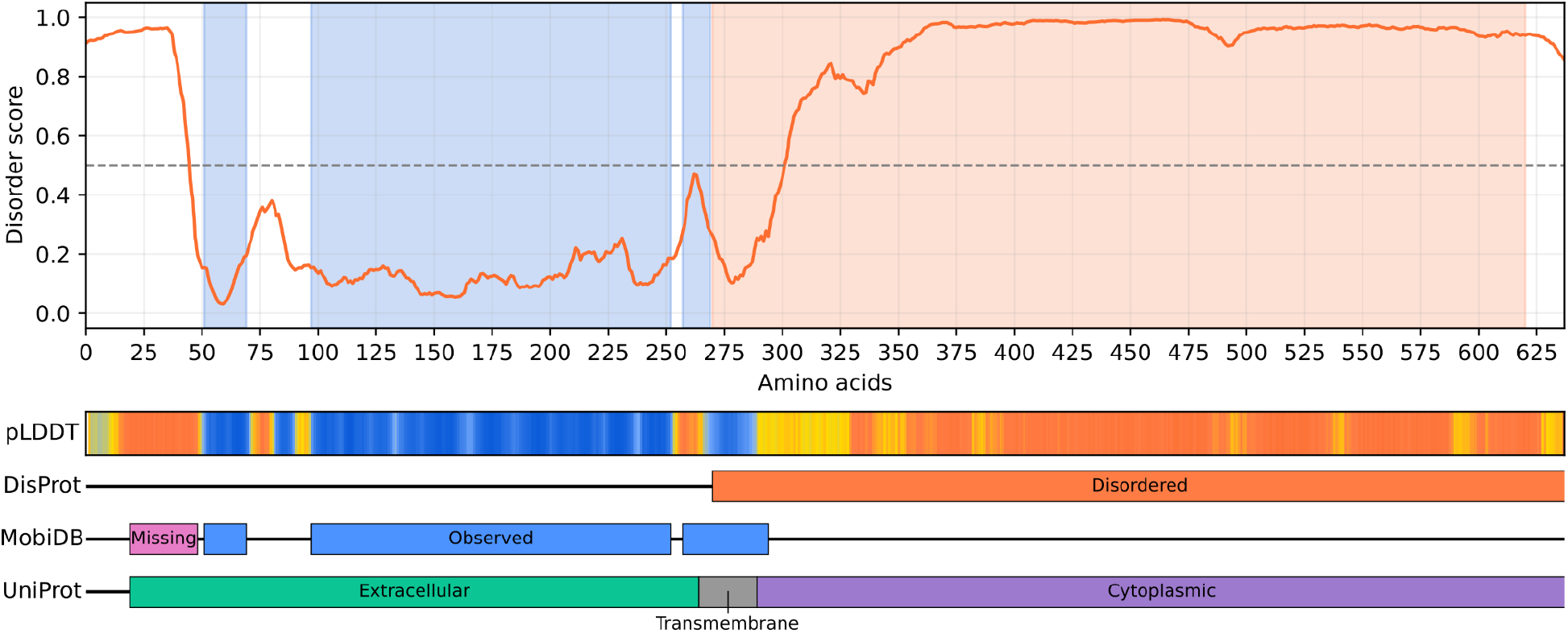
Disorder prediction for the training protein P10912 (DisProt ID: DP00033). The curve shows the emb2dis score for disorder at each position, while the ground-truth annotations from DisProt are shown as shaded regions. Bottom: AF2 pLDDT score, DisProt, MobiDB, and UniProt annotations.

Figure 4 shows an example of emb2dis-ESM2 prediction for the protein Q8GXC2 (DisProt ID: DP04240), Myb family transcription factor PHL4, which is in the CAID3 challenge test set. This transcription factor is involved in male gametophyte development and is essential for plant response to low levels of phosphate. Proteins in this transcription factor family act by altering various gene expression levels, such as increasing levels of the acid phosphatase proteins, which catalyze the conversion of inorganic phosphates to bio-available compounds (Fonda & Murray, 2024). Q8GXC2 protein has two portions of annotated disorder regions (1-227 and 369-397), which are perfectly detected by emb2dis, and this also largely coincides with the low AF2 pLDDT score. Interestingly, emb2dis also indicates the region 291-319 as disordered with a high score, in accordance with the low pLDDT scores. In spite of not being labeled as disorder, this region is between two annotated ordered regions, a Myb-like DNA-binding domain (PF00249) (233-284) and a Coiled coil domain (319-339), and it might be the discovery of a new disorder annotation region.

**Figure 4.**
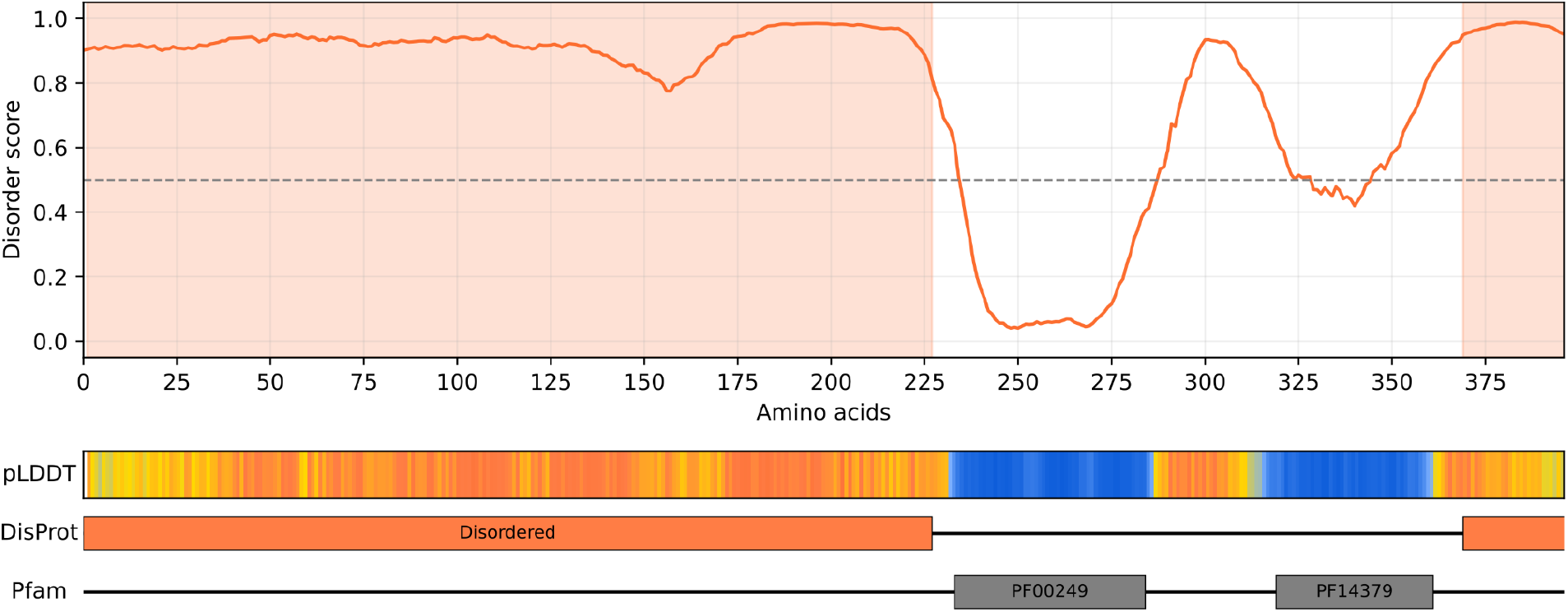
Disorder prediction for the protein Q8GXC2 (DisProt ID: DP04240), of the CAID3 challenge test set. The curve shows the disorder score at each position, while the ground-truth annotations from DisProt are shown as shaded regions. Bottom: AF2 pLDDT score, DisProt, and Pfam annotations.

Finally, Figure 5 shows another test protein, Q8N6T7 (DisProt ID: DP04167), a NAD^+^-dependent histone deacylase, known as Sirtuin-6, which plays an essential role in DNA damage repair, telomere maintenance, metabolic homeostasis, inflammation, tumorigenesis and aging (Michishita et al, 2008). This is a very interesting case in which our model, emb2dis-ESM2, has correctly detected all three IDRs. However, AlphaFold2 assigns a high confidence score to residues 62-84, despite being annotated in DisProt and reported in literature as disordered. Rather than indicating a fully ordered region, these high pLDDT values may suggest that this segment can adopt a more defined conformation under specific conditions, known as context-dependent folding (Piovesan et al., 2022, Alderson et al., 2023). Within this region, MobiDB annotates observed residues. Nevertheless, the correct identification of these residues as disordered by emb2dis-ESM2 indicates that this model is able to identify disorder in regions undergoing context-dependent folding, while AF2 cannot. This is a remarkable result that highlights the practical utility of emb2dis for disorder prediction, as a reliable and accurate tool in comparison to more costly computational methods.

**Figure 5.**
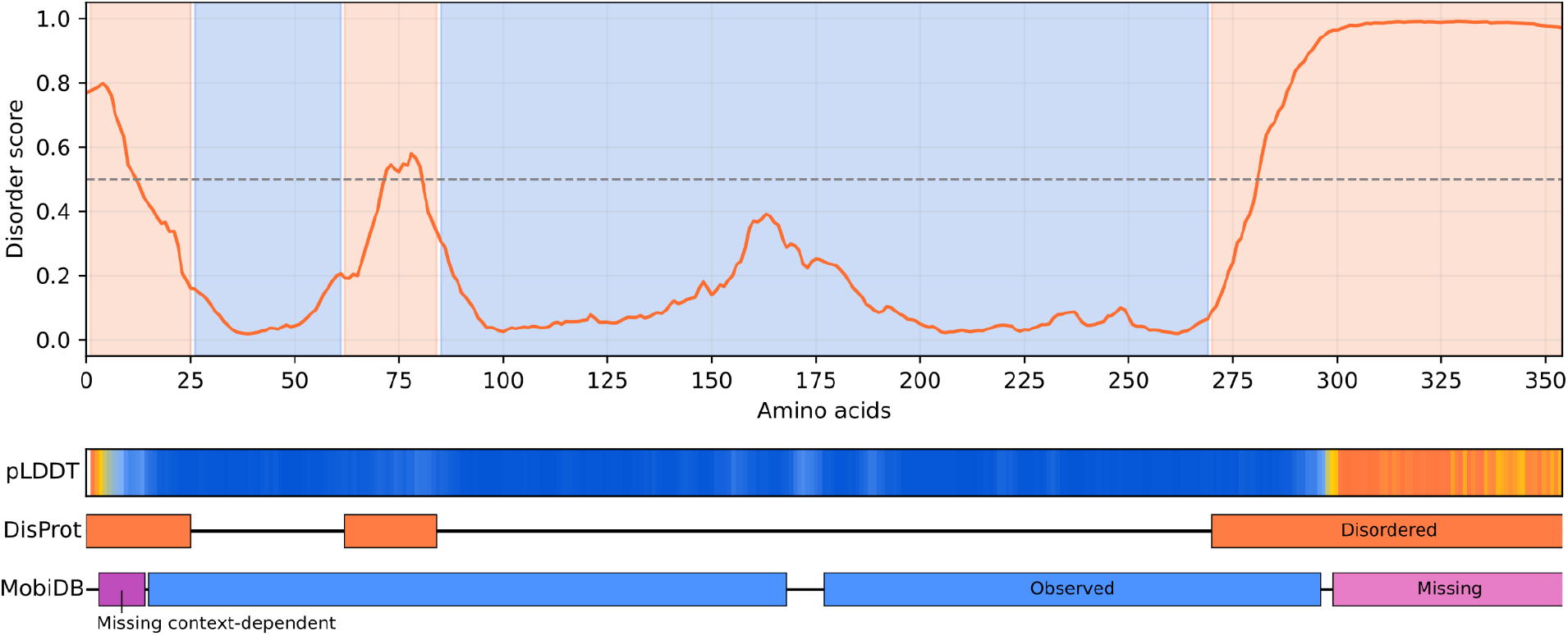
Disorder prediction for the protein Q8N6T7 (DisProt ID: DP04167), of the CAID3 challenge test set. The curve shows the predicted disorder score at each position, while the ground-truth annotations from DisProt are shown as shaded regions. Bottom: AF2 pLDDT score, DisProt, and MobiDB annotations.

## Conclusions

In this work we introduced emb2dis, a computational model that demonstrated high effectiveness for predicting disorder from protein sequences, assigning a disorder propensity score to each residue in the sequence. Differently to existing state-of-the-art approaches for this task, which rely on pLM representations and standard convolutional layers, the success of emb2dis highlights the advantages of the novel DL architecture that combines residual networks with dilated convolutions. Since dilated convolutions increase the receptive field of convolutions, the model can capture more effectively the local and global context of each amino acid, which is particularly useful for detecting disordered regions. The emb2dis model was evaluated on the latest CAID3 blind benchmark datasets for disorder prediction, with three different pLMs. The emb2dis-ESM2 model showed the best results, achieving first place in the Disorder-PDB and ranking top-10 in the Disorder-NOX. Future work will focus on improving the model predictions for this last dataset, in particular by refining the labeling of training data.

## Availability and usage

The emb2dis predictor is freely available and it can be found in https://sinc.unl.edu.ar/web-demo/emb2dis/, where input sequences of up to 1,000 amino acids can be processed online with this web-demo. To process longer sequences emb2dis can be installed from the repository https://github.com/sinc-lab/emb2dis. In the input interface of the emb2dis online tool, the user can enter a protein sequence, or load a .fasta file with the sequence. The language model to be used for the protein representation can be selected from a list of available pre-trained models. The emb2dis output shows the total length of the proteín analyzed, the number of disorder residues and the disorder percentage of the protein. The predicted disorder region is indicated with a color and the curve of the predictor is shown as well. The detailed prediction of disorder for the input proteins, per amino acid, can be downloaded in a .csv file.

## Acknowledgments

This publication is part of a project that has received funding from HORIZON-MSCA-SE project IDPfun2 (no. 101182949)

